# I^3^S Classic and Insect Species Identification of Diptera and Hymenoptera (Mosquitoes and Bees)

**DOI:** 10.1101/090621

**Authors:** Nayna Vyas-Patel, John D Mumford

## Abstract

A number of image recognition systems have been specifically formulated for the individual recognition of large animals. These programs are versatile and can easily be adapted for the identification of smaller individuals such as insects. The Interactive Individual Identification System, I^3^S Classic, initially produced for the identification of individual whale sharks was employed to distinguish between different species of mosquitoes and bees, utilising the distinctive vein pattern present on insect wings. I^3^S Classic proved to be highly effective and accurate in identifying different species and sexes of mosquitoes and bees, with 80% to100% accuracy for the majority of the species tested. The sibling species *Apis mellifera* and *Apis mellifera carnica* were both identified with100% accuracy. *Bombus terrestris terrestris* and *Bombus terrestris audax*; were also identified and separated with high degrees of accuracy (90% to 100% respectively for the fore wings and 100% for the hind wings). When both *Anopheles gambiae* sensu stricto and *Anopheles arabiensis* were present in the database, they were identified with 94% and 100% accuracy respectively, allowing for a morphological and non-molecular method of sorting between these members of the sibling complex. Flat, not folded and entire, rather than broken, wing specimens were required for accurate identification. Only one wing image of each sex was required in the database to retrieve high levels of accurate results in the majority of species tested. The study describes how I^3^S was used to identify different insect species and draws comparisons with the use of the CO1 algorithm. As with CO1, I^3^S Classic proved to be suitable software which could reliably be used to aid the accurate identification of insect species. It is emphasised that image recognition for insect species should always be used in conjunction with other identifying characters in addition to the wings, as is the norm when identifying species using traditional taxonomic keys.

## Introduction

The need to develop non-invasive methods of recognising individual animals in the field, led to the development of different image recognition systems which are widely used today, by both the professional and the citizen scientist (Moro, 2014). To avoid having to capture/recapture and mark/tag animals from the wild, photos were used for individual animal identification and landmark features and markings on the body noted and compared. However, this required the investigator to physically go through a large number of previously stored photographs in order to compare, match and identify images of individuals at different times. This was extremely time consuming, prone to human error and not always accurate and image recognition software began to be employed to take on this arduous task (Bolgar et al, 2012; Dillow, 2010; Knight, 2010; Speed et al, 2007). These systems were initially developed with specific animals in mind, however, the software could just as easily be applied to the recognition of other species. One such method is the freely available Interactive Individual Identification system, I^3^S, initially developed for the recognition of individual marine animals such as whale sharks and manta rays (Hartog and Reijns, 2013).

The venation present on insect wings is unique for a given species and this distinctive ‘fingerprint’ is often used as one of the reliable features for insect species identification. Another advantage of using dead insect wings for species identification is that unlike a large, living animal, it is not a moving object, allowing time to obtain the best possible image, this helps to reduce any image related errors as outlined by Bolgar et al (2012), who used ‘Wild ID’ to identify giraffes. In this study, I^3^S was utilised to identify different species of Dipteran and Hymenopteran insects, using images of their wings.

I^3^S utilises the pattern of spots on an animal to determine identification. Whilst the insect species investigated here did not possess spots on their wings, the points of intersection of the veins and where the veins met the periphery of the wing could reliably be used as unique ‘markers’ for the species. A number of semi-automated systems utilise insect wing veins to identify species, a system known as ‘Geometric Morphometrics’ (Dujardin, 2011; Wilke ABB, 2016). Francoy et al (2008), described two fast and efficient procedures based on geometric morphometrics and the bee identification program ABIS (Automatic Bee Identification system). Like ABIS, ‘DrawWing’ is semi-automated software designed primarily for the identification of bee species (Tofilski, 2004). A study by Lorenz et al (2012) outlined how geometric morphometrics was successfully used to distinguish between three Anopheline mosquito vectors from Brazil. Some methods utilised an entire sketch of the wing, or the area bounded by the veins, ‘the cells’, whilst others used the junctions and intersections of veins to identify species (Hall, 2011). However, whilst the wing features and markings used in geometric morphometric methods were excellent, they did not benefit from the advanced features of image recognition software available at the current time, such as the possibility of creating and using newly formed databases capable of storing and retrieving accurate matches from thousands of stored images. In the present study, the points of intersection of the veins both within the wing and at the periphery of the wing were used as markers in I^3^S and the accuracy of the software in identifying the species was assessed.

I^3^S presents a choice of different software options – Classic, Pattern, Contour and Spot. On the recommendation of the creators of the software (Hartog and Reijns, 2013), I^3^S Classic was utilised for the identification of different species of mosquitoes, bees and bumblebees.

As with other image recognition software, I^3^S ranks the identified results, so that the closest match from the database is ranked 1 and the next closest is ranked thereafter according to how closely the marked area resembles the test image. A score is automatically assigned to each match and all of the ranked results can be seen by eye. For any given identification, if the result at rank 1 did not match the test species as seen by eye, or the score was far removed from 0 (a score of 0 being an exact copy), it could be queried and examined further.

The number of images of each species which are required in the database for optimum species recognition was examined. A previous study using different software, where the whole image was examined without marking salient features on the image prior to testing, indicated that databases containing larger numbers of each species produced higher numbers of accurate results than using databases containing smaller numbers of each species (Vyas-Patel et al, 2015). Different sizes of databases were therefore created and analysed for their accuracy in identifying the species of a test image.

Initially, using a ‘large database’ (LDB) containing 20 images of each species (10 of each sex) was compared to using a ‘small database’ (SDB) containing 10 images of each species (five of each sex) to determine if a using a larger database resulted in greater species identification accuracy than using a smaller database.

The minimum number of images of each species required in the database to produce high numbers of accurate identifications was explored. To begin with just one image of each species was inputted into the database, tested with new images of different species and the results noted. This was followed by using two images (one of each sex) of each species in the database.

The aim of creating differently sized databases was to determine the levels of accurately identified species in rank 1 from databases containing just one image, two images, 10 images and 20 representative images of each of the species to be tested. The numbers of accurately identified species from rank 1 to 5 and 1 to 10 was also examined when there were at least 5 or 10 images of each species being tested in the database.

An earlier study using different software noted that if there was more than one image in the database of a particular species, when tested with a wing of that species, all of the correct images from the database appeared consecutively correctly in the ranked results (Vyas-Patel et al, 2015). If there were five images of a species in the database, when tested with the same species, all five images from the database were retrieved from rank 1 to rank 5. This added to the certainty that the correct species had been identified, provided sufficient images of the test were present in the database. This consecutive ranking of the correct species was therefore examined here.

To ascertain if using different models of cameras affected the results, different cameras were used to take photos of insect wings and the results compared. Finally the study explored the use of I^3^S in identifying 38 different species of mosquito and bee wings donated from different places, both wild caught and laboratory reared.

## Method

### Preparation and marking of the Images

Insect specimens were obtained from a wide variety of donors and locations as outlined in Table 1. The wings were dissected from the body under a standard dissection microscope and photographed with a Samsung NV10 digital camera, using only the sub-stage lighting of the microscope, as this produced a clear image of the wing shape and venation. Prior to taking a photograph, a microscope cover slip was placed on the wing to ensure that any folds were gently flattened out. This was not necessary for the Hymenopteran (bee and bumblebee) wings. Each image was uploaded into an Adobe Photoshop (CS5) image editor and rotated so that the point of insertion of the wing into the body of the insect always faced to the left and the wing was aligned to be as horizontal as possible, using Image Rotate in the top menu bar of Photoshop. The newly aligned and rotated images were saved as .jpg files, creating a different file for each species, sex and where appropriate, fore and hind wings.

**Table 1:**
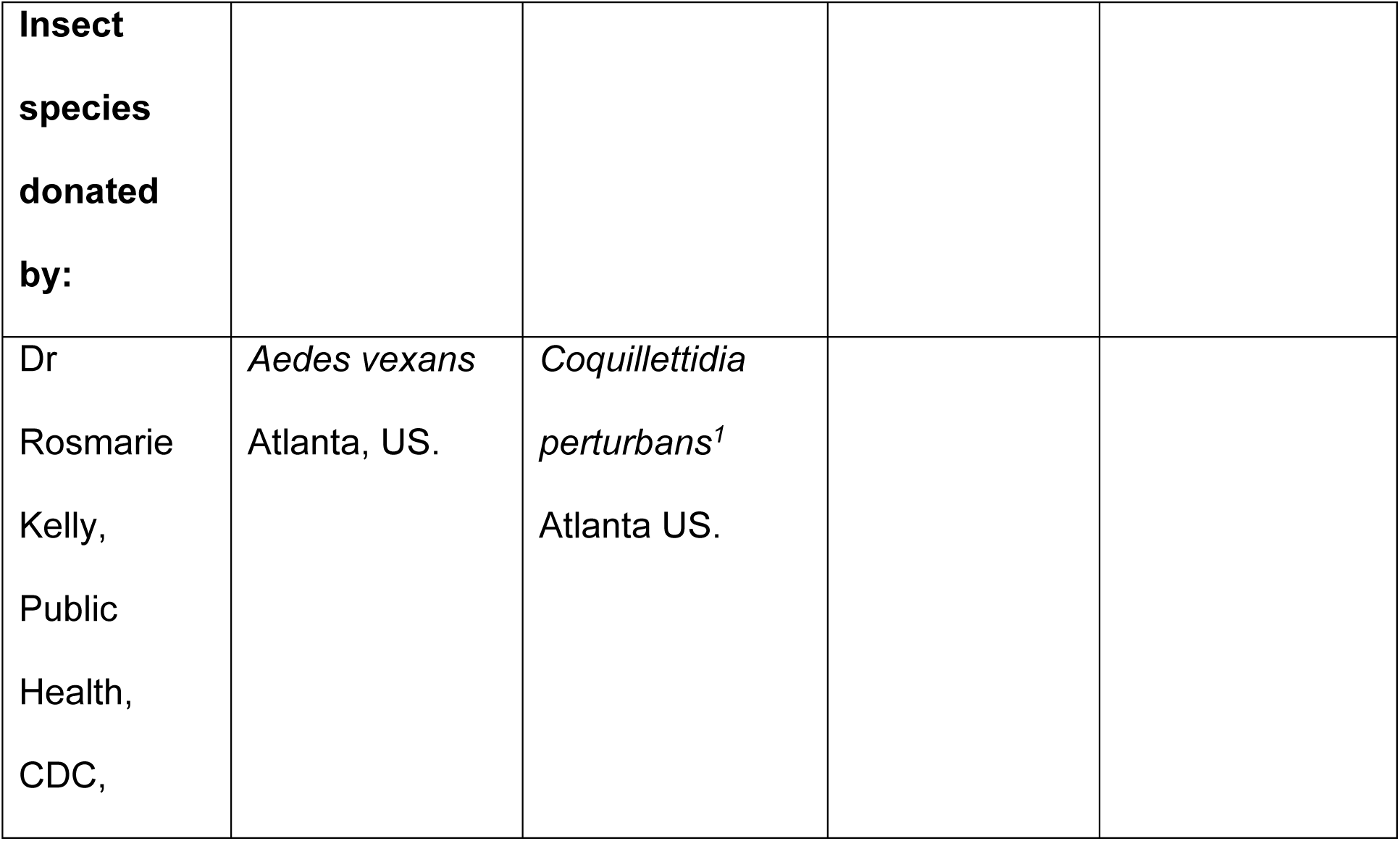
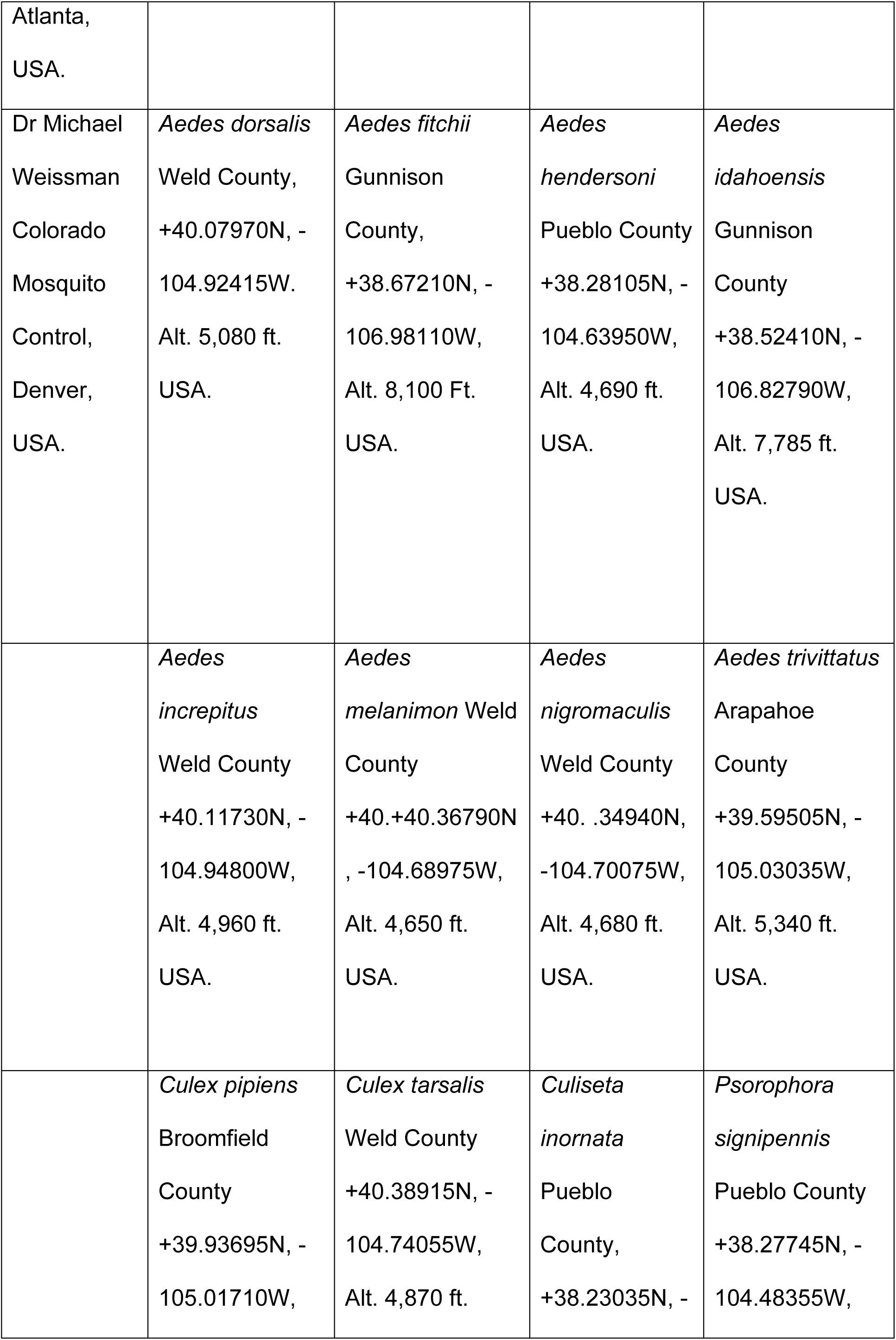
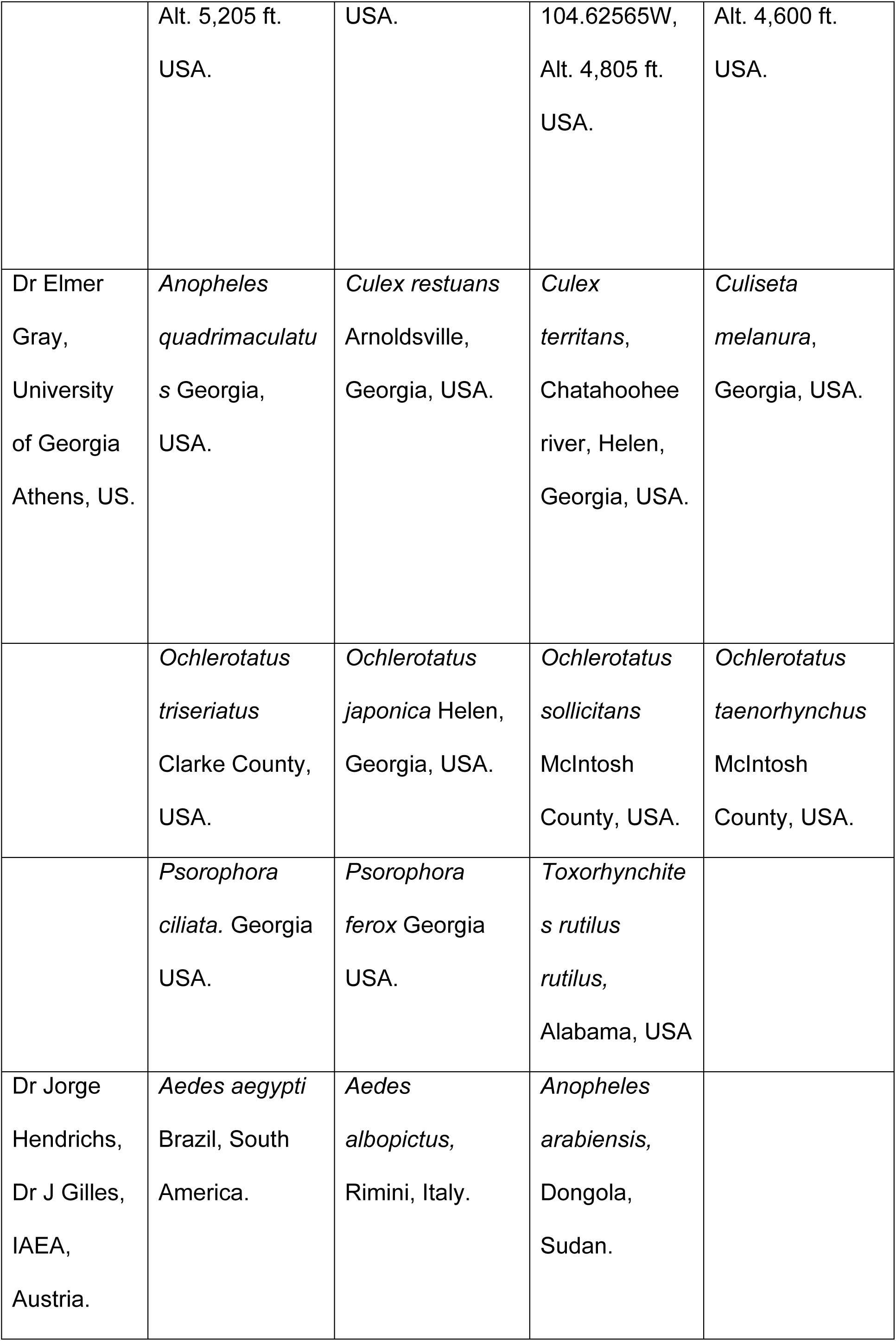
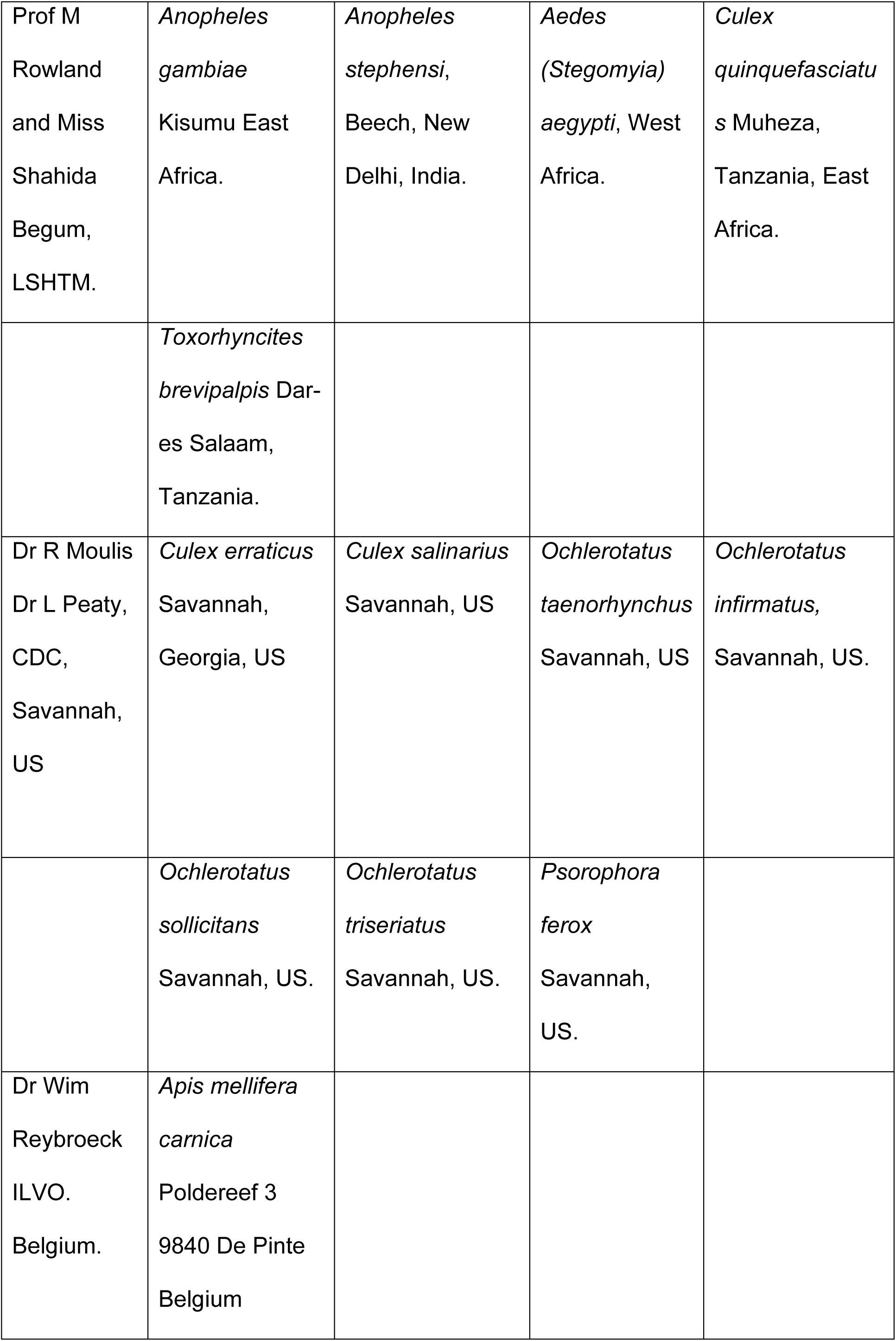
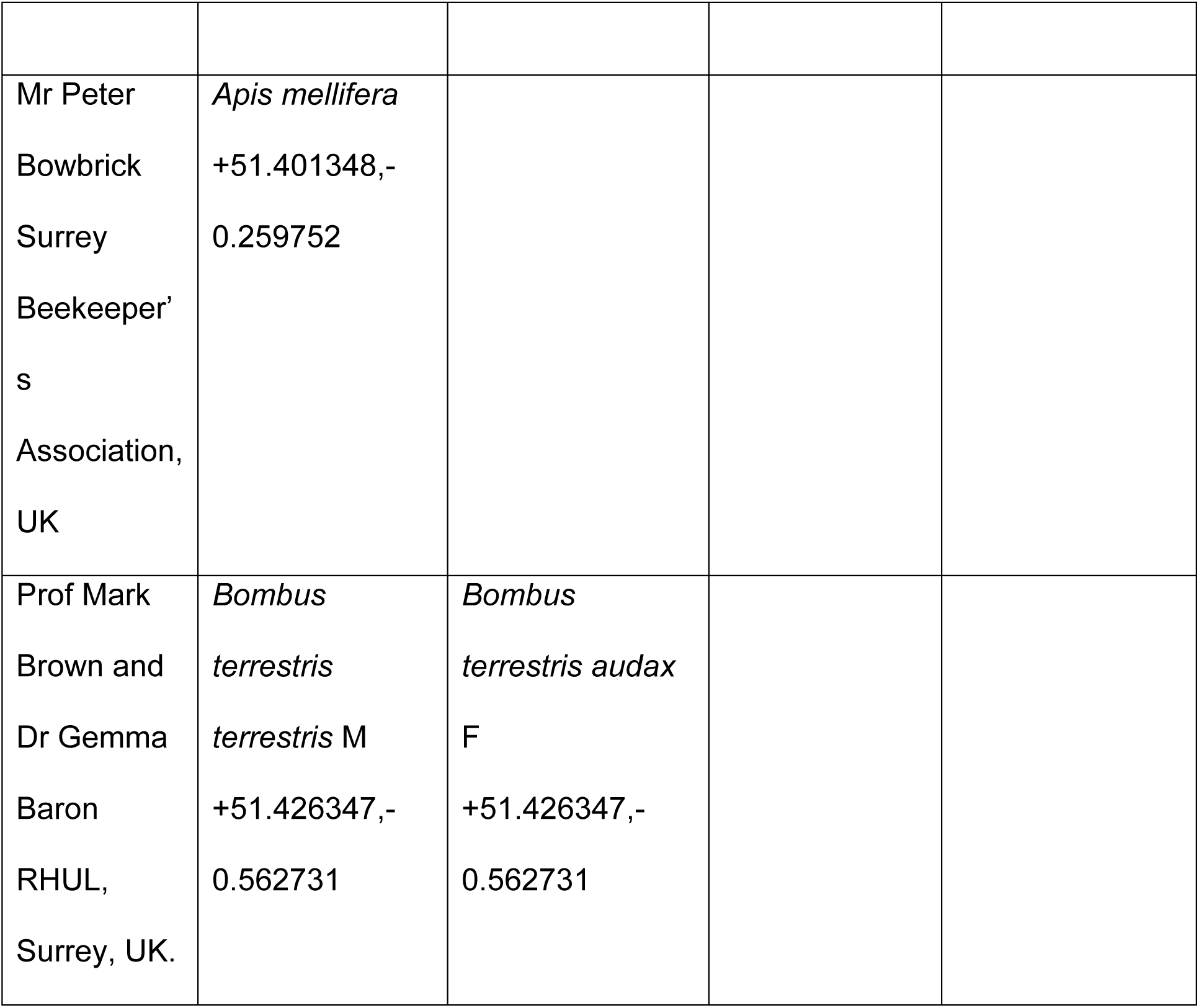
Insect Donors & Available Location Data for the different Mosquito, Bee & Bumblebee Species Used.

| **Insect species donated by:** |  |  |
| --- | --- | --- |
| Dr<br>Rosmarie<br>Kelly,<br>Public<br>Health,<br>CDC, | *Aedes vexans*<br>Atlanta, US. | *Coquillettidia*<br>*perturbans*<sup>1</sup><br>Atlanta US. |

|  |  |  |  |  |
| --- | --- | --- | --- | --- |
| Atlanta,<br>USA. |  |  |  |  |
| Dr Michael<br>Weissman<br>Colorado<br>Mosquito<br>Control,<br>Denver,<br>USA. | *Aedes dorsalis*<br>Weld County,<br>+40.07970N, -<br>104.92415W.<br>Alt. 5,080 ft.<br>USA. | *Aedes fitchii*<br>Gunnison<br>County,<br>+38.67210N, -<br>106.98110W,<br>Alt. 8,100 Ft.<br>USA. | *Aedes*<br>*hendersoni*<br>Pueblo County<br>+38.28105N, -<br>104.63950W,<br>Alt. 4,690 ft.<br>USA. | *Aedes*<br>*idahoensis*<br>Gunnison<br>County<br>+38.52410N, -<br>106.82790W,<br>Alt. 7,785 ft.<br>USA. |
|  | *Aedes*<br>*increpitus*<br>Weld County<br>+40.11730N, -<br>104.94800W,<br>Alt. 4,960 ft.<br>USA. | *Aedes*<br>*melanimon* Weld<br>County<br>+40.+40.36790N<br>, -104.68975W,<br>Alt. 4,650 ft.<br>USA. | *Aedes*<br>*nigromaculis*<br>Weld County<br>+40. .34940N,<br>-104.70075W,<br>Alt. 4,680 ft.<br>USA. | *Aedes trivittatus*<br>Arapahoe<br>County<br>+39.59505N, -<br>105.03035W,<br>Alt. 5,340 ft.<br>USA. |
|  | *Culex pipiens*<br>Broomfield<br>County<br>+39.93695N, -<br>105.01710W, | *Culex tarsalis*<br>Weld County<br>+40.38915N, -<br>104.74055W,<br>Alt. 4,870 ft. | *Culiseta*<br>*inornata*<br>Pueblo<br>County,<br>+38.23035N, - | *Psorophora*<br>*signipennis*<br>Pueblo County<br>+38.27745N, -<br>104.48355W, |

|  |  |  |  |  |
| --- | --- | --- | --- | --- |
|  | Alt. 5,205 ft.<br>USA. | USA. | 104.62565W,<br>Alt. 4,805 ft.<br>USA. | Alt. 4,600 ft.<br>USA. |
| Dr Elmer<br>Gray,<br>University<br>of Georgia<br>Athens, US. | *Anopheles*<br>*quadrimaculatus*<br>Georgia,<br>USA. | *Culex restuans*<br>Arnoldsville,<br>Georgia, USA. | *Culex*<br>*territans*,<br>Chatahoochee<br>river, Helen,<br>Georgia, USA. | *Culiseta*<br>*melanura*,<br>Georgia, USA. |
|  | *Ochlerotatus*<br>*triseriatus*<br>Clarke County,<br>USA. | *Ochlerotatus*<br>*japonica* Helen,<br>Georgia, USA. | *Ochlerotatus*<br>*sollicitans*<br>McIntosh<br>County, USA. | *Ochlerotatus*<br>*taeniorhynchus*<br>McIntosh<br>County, USA. |
|  | *Psorophora*<br>*ciliata*. Georgia<br>USA. | *Psorophora*<br>*ferox* Georgia<br>USA. | *Toxorhynchites*<br>*rutilus*<br>*rutilus*,<br>Alabama, USA |  |
| Dr Jorge<br>Hendrichs,<br>Dr J Gilles,<br>IAEA,<br>Austria. | *Aedes aegypti*<br>Brazil, South<br>America. | *Aedes*<br>*albopictus*,<br>Rimini, Italy. | *Anopheles*<br>*arabiensis*,<br>Dongola,<br>Sudan. |  |

|  |  |  |  |  |
| --- | --- | --- | --- | --- |
| Prof M Rowland and Miss Shahida Begum, LSHTM. | *Anopheles gambiae*<br>Kisumu East Africa. | *Anopheles stephensi*,<br>Beech, New Delhi, India. | *Aedes (Stegomyia) aegypti*, West Africa. | *Culex quinquefasciatus* Muheza, Tanzania, East Africa. |
|  | *Toxorhynchites brevipalpis* Dar-es Salaam, Tanzania. |  |  |  |
| Dr R Moulis<br>Dr L Peaty, CDC, Savannah, US | *Culex erraticus*<br>Savannah, Georgia, US | *Culex salinarius*<br>Savannah, US | *Ochlerotatus taeniorhynchus*<br>Savannah, US | *Ochlerotatus infirmatus*,<br>Savannah, US. |
|  | *Ochlerotatus sollicitans*<br>Savannah, US. | *Ochlerotatus triseriatus*<br>Savannah, US. | *Psorophora ferox*<br>Savannah, US. |  |
| Dr Wim Reybroeck ILVO. Belgium. | *Apis mellifera carnica*<br>Poldereef 3<br>9840 De Pinte<br>Belgium |  |  |  |
| Mr Peter Bowbrick<br>Surrey<br>Beekeeper's<br>Association,<br>UK | *Apis mellifera*<br>+51.401348,-<br>0.259752 |  |
| Prof Mark Brown and<br>Dr Gemma Baron<br>RHUL,<br>Surrey, UK. | *Bombus terrestris*<br>*terrestris* M<br>+51.426347,-<br>0.562731 | *Bombus terrestris audax*<br>F<br>+51.426347,-<br>0.562731 |

I^3^S Classic, version 4.01, was downloaded from the internet and a ‘fingerprint’ (.fpg) image of each wing image was prepared and stored in the database as shown in Figure 2. The first three reference points (in blue, made larger for ease of viewing) were selected as shown in Figure 2. I^3^S allows these 3 points in blue to be renamed within the program – this will not affect the results. Further information on the placement of the three blue reference points is given in the frequently asked questions (FAQs) on the I^3^S website. This was followed by marking the points of intersection of the veins within the wing and where the veins actually met the edge or their extrapolation to the edge of the wing (red points in Figure 2). Any clearly visible point and landmark wing feature may be chosen for marking as long as subsequent images are also consistently marked in the same way for a given species. The default setting of I^3^S requires a minimum of 12 and a maximum of 30 red reference points to be marked out. The numbers of maximum and minimum reference points in red can be changed within the program, but was kept to the default setting for this study. Where the minimum was not possible, as in the case of bee hind wings, points near the edge of the wings were extrapolated to meet the edge of the wing as shown in Figure 2. Each fingerprint file (.fgp) was saved and could then be used either for database creation or as a test image. A comprehensive guide to the use of I^3^S and how images can be prepared, stored and analysed is given in the instructions on the I^3^S website together with a tutorial.

**Figure 2:**
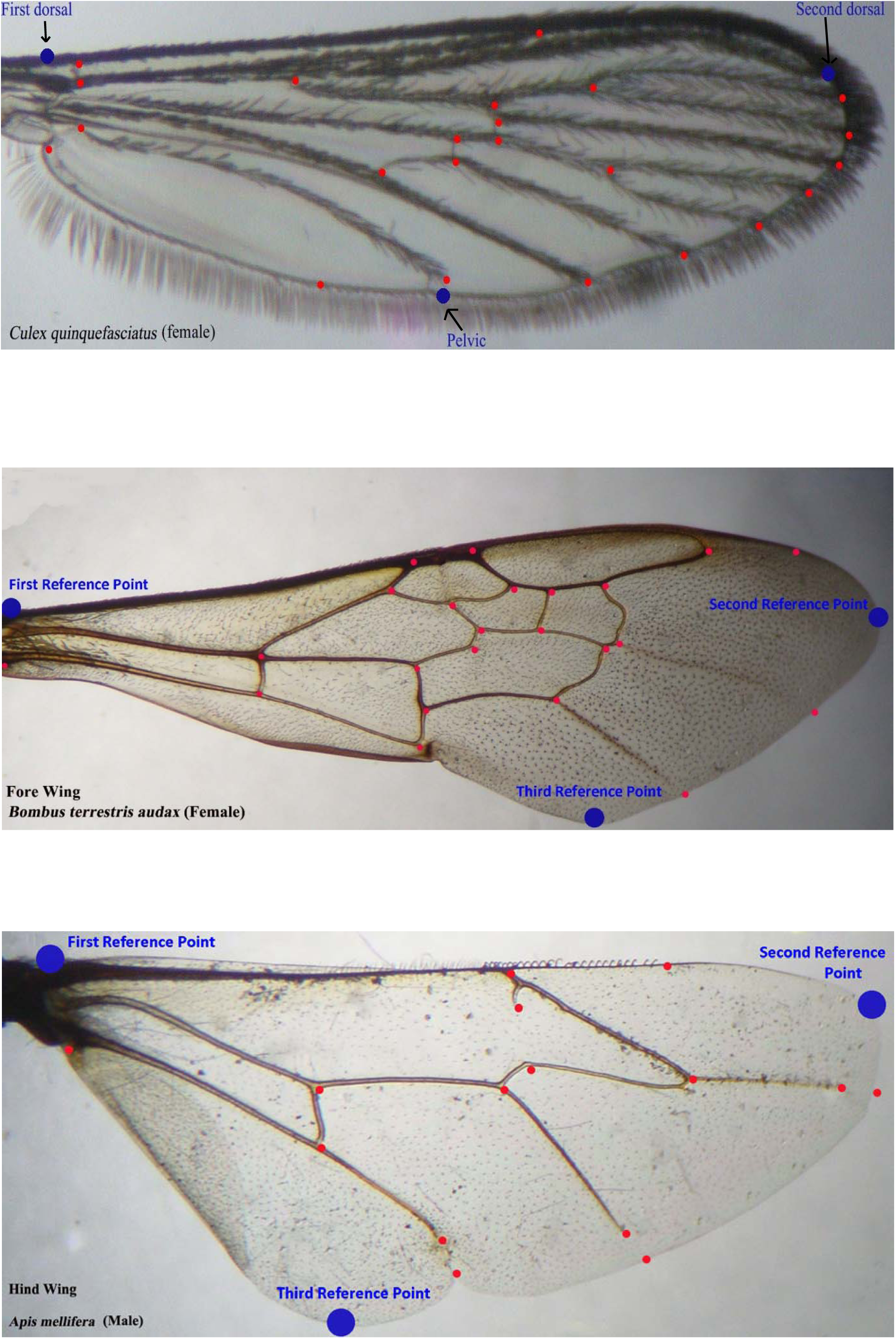
Indicating the Points of Intersection of the Veins Marked on Sample Wing Images.

### Database creation

A large database (LDB) of fingerprint .fgp, files was created. This comprised of 20 images each (10 of each sex) of *Anopheles gambiae, Anopheles stephensi, Aedes aegypti, Culex quinquefasciatus and Toxorhynchites brevipalpis* mosquito species; similarly, 20 images each of *Apis mellifera* honey bees and the two bumblebee subspecies. In the case of the Hymenopteran species, 20 images of the fore and 20 images of the hind wings of each species were uploaded onto the large database (mosquitoes do not possess hind wings, these being reduced to small stumps called halteres). Only one sex was available of the *Bombus* subspecies, *Bombus terrestris terrestris* (Btt) and *Bombus terrestris audax* (Bta) and 10 images each of the fore and hind wings of each subspecies were uploaded into the database. In the case of *A. mellifera* both sexes were available and 10 each of the sexes of fore and hind wings were used (40 images in total for *A. mellifera*). The large database therefore contained a total of 180 different images of both Dipteran and Hymenopteran wing images.

A different, smaller, database was also created (SDB), this time containing 10 images of each species comprising of five males and five females, therefore 10 images each of the five mosquito species making a total of 50 mosquito wing images. In *Apis mellifera* where both sexes were available, five fore and five hind wings of both sexes were used, total 20 images were added to the database. In the bumble bee species, five fore and five hind wings of Btt (total 10) and Bta (total 10) each were uploaded. A grand total of 90 images of the different species were present in the smaller database or SDB.

A third database (1-DB) was created with only one representative image of each species of the Diptera and Hymenoptera above, this included one image each of the hind and fore wings of the Hymenoptera. A grand total of 11 images were present in the database containing only one example wing image of each of the species. A fourth database (2-DB) was created with just 2 images, one of each sex from each of the species to be tested.

Fifty mosquito and bee of each species and 35 bumblebee images of each sub species were tested using each of the databases. The percentage of correctly identified results at rank 1 was noted for all of the different databases created, Table 2. The numbers of correctly identified species within rank 1 to 5 and rank 1 to 10 were totalled for each test image and the most frequently occurring total (the mode) was noted, Table 2a.

**Table 2:** Total percentages of correctly identified species at rank 1, I^3^S Classic, using different Databases.

|  | 1-DB %<br>Rank 1<br>Species<br>Correct | LDB %<br>Rank 1<br>Species<br>Correct | SDB<br>% Rank 1<br>Species<br>Correct | 2-DB<br>% Rank 1<br>Species<br>Correct |
| --- | --- | --- | --- | --- |
| *A. Stephensi* | 60% | 100% | 100% | 100% |
| *A. gambiae* | 57% | 100% | 100% | 100% |
| *A. aegypti* | 93% | 100% | 100% | 100% |
| *C. quinquefasciatus* | 58% | 100% | 100% | 100% |
| *T. brevipalpis* | 97% | 100% | 100% | 100% |
| *A. mellifera* Fore<br>Wings | 100% | 100% | 100% | 100% |
| *A. mellifera* Hind<br>Wings | 100% | 100% | 100% | 100% |
| *B. terrestris*<br>*terrestris* (Btt) Fore<br>Wings | 75% | 97% | 87% | N/A |
| *B. terrestris audax*<br>(Bta) Fore Wings | 40% | 100% | 83% | N/A |
| Btt Hind Wings | 56% | 93% | 93% | N/A |
| Bta Hind Wings | 67% | 100% | 100% | N/A |
| LDB, SDB t value |  |  | 1.2394 | N/A |
| LDB, SDB p value |  |  | 0.2296 | N/A |
| 1-DB, LDB t value |  | **4.018\*** |  | N/A |
| 1-DB, LDB p value |  | **0.0007\*** |  | N/A |
| 1-DB, SDB t value | **3.547\*** |  |  | N/A |
| 1-DB, SDB p value | **0.002\*** |  |  | N/A |

**Asterisk \* indicates statistically significant results at $p < 0.05$ .**

**Table 2a:** The Modal Values Up to Rank 5 and Rank 10, from the Larger (LDB) & Smaller (SDB) Databases, I^3^S Classic.

|  | LDB<br>Modes<br>Rank 5 | SDB<br>Modes<br>Rank 5 | LDB<br>Modes<br>Rank 10 | SDB<br>Modes<br>Rank 10 |
| --- | --- | --- | --- | --- |
| *A. Stephensi* | 5 | 5 | 9 | 7 |
| *A. gambiae* | 5 | 5 | 9 | 7 |
| *A. aegypti* | 5 | 5 | 9 | 8 |
| *C. quinquefasciatus* | 5 | 5 | 9 | 8 |
| *T. brevipalpis* | 5 | 5 | 10 | 7 |
| *A. mellifera* Fore<br>Wings | 5 | 5 | 10 | 10 |
| *A. mellifera* Hind<br>Wings | 5 | 5 | 10 | 10 |
| *B. terrestris*<br>*terrestris* (Btt) Fore<br>Wings | 4 | 4 | 8 | 6 |
| *B. terrestris audax* | 5 | 3 | 10 | 7 |
| (Bta) Fore Wings |  |  |  |  |
| Btt Hind Wings | 5 | 3 | 9 | 7 |
| Bta Hind Wings | 5 | 4 | 9 | 6 |
| t value |  | 1.7252 |  | **3.7849\*** |
| p value |  | 0.099 |  | **0.0012\*** |

**Asterisk \* indicates statistically significant results at $p < 0.05$ .**

### Consecutively Correct Results

Five wing images of each sex from five different mosquito species and five fore and five hind wings of *A. mellifera* were uploaded into the database and tested with a total of 30 test images. The results of each test were noted up to rank 5 and the total numbers of consecutively and correctly identified species noted, Table 3 and Figure 1.

**Figure 1:**
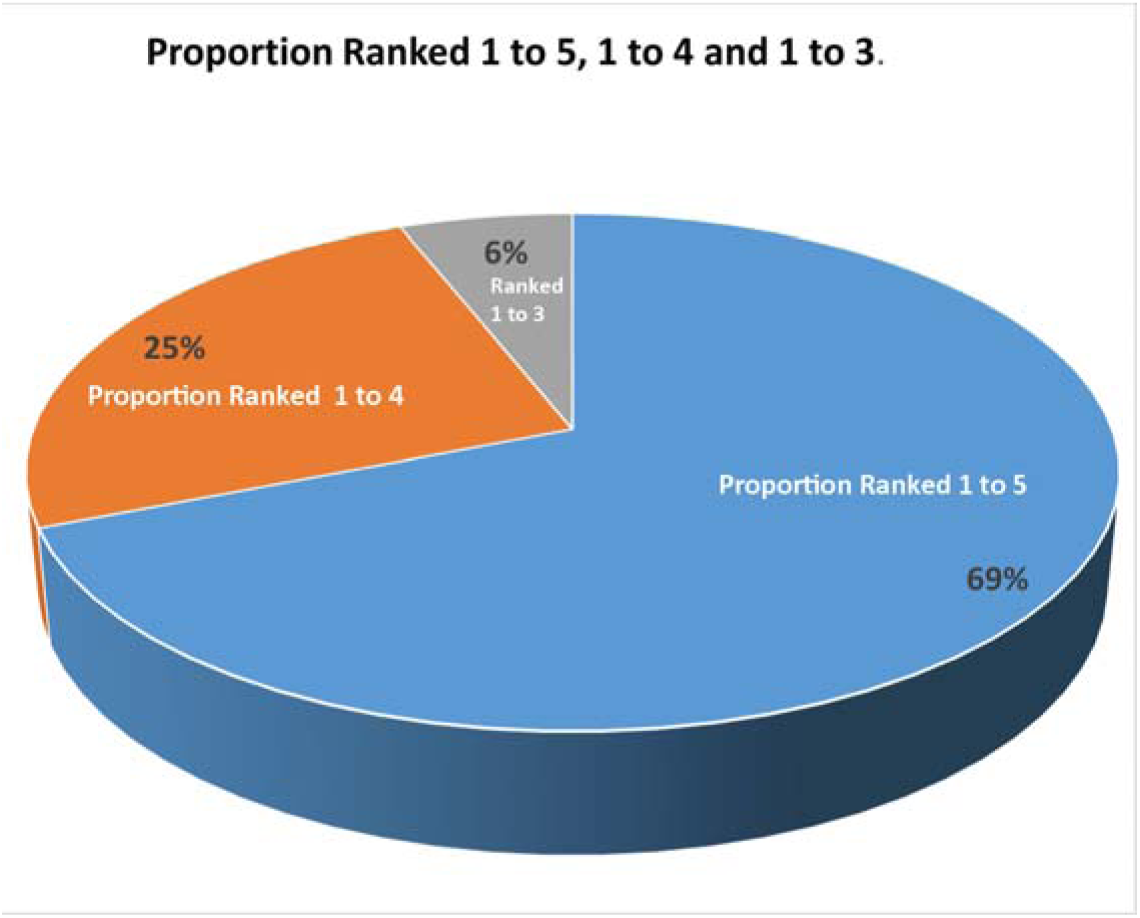
Indicating the percentage of species correctly and consecutively ranked from rank 1 to 5, 1 to 4 and 1 to 3.

**Table 3:** The percentage of consecutively & correctly identified species when five wing images of each sex (total 10 of each species) were present in the Database.

|  |  |  |  |
| --- | --- | --- | --- |
|  | Total % with<br>the first 5 ranks<br>consecutively | Total % with<br>the first 4 ranks<br>consecutively | Total % with<br>the first 3 ranks<br>consecutively |
|  | correct | correct | correct |
| *Anopheles gambiae* F | 35% | 40% | 25% |
| *Anopheles gambiae* M | 80% | 20% |  |
| *Anopheles stephensi* F | 40% | 40% | 20% |
| *Anopheles stephensi* M | 80% | 20% |  |
| *Aedes aegypti* F | 68% | 32% |  |
| *Aedes aegypti* M | 69% | 31% |  |
| *Culex quinquefasciatus* F | 55% | 45% |  |
| *Culex quinquefasciatus* M | 29% | 30% | 41% |
| *Toxorhyncites brevipalpis* F | 85% | 15% |  |
| *Toxorhyncites brevipalpis* M | 66% | 34% |  |
| *Apis mellifera* Fore F | 70% | 30% |  |
| *Apis mellifera*<br>Fore M | 90% | 10% |  |
| *Apis mellifera*<br>Hind F | 100% |  |  |
| *Apis mellifera*<br>Hind M | 100% |  |  |
| Grand Total %<br>in each<br>category* | 69% | 25% | 6% |

\* = To ascertain the grand total % of correctly identified species up to rank 5, 4 and 3, the totals were added in each category (column), divided by the grand total and multiplied by 100. There were no cases where only the first two ranks were correctly identified. They were all correctly and consecutively identified at least up to rank 3 as depicted in the associated Figure 1 below:

### Models of camera

Photographs of five different species of mosquitoes were taken with different models of cameras and the results noted for rank 1 identifications (Table 4). The database used photos taken with the Samsung NV10; five images each of male and female wings of *Anopheles gambiae, Anopheles stephensi, Aedes aegypti, Culex quinquefasciatus* and *Toxorhyncites brevipalpis* were uploaded into the database and tested with 30 images (15 of each sex) from each of the different models of cameras (Table 4).

**Table 4:**
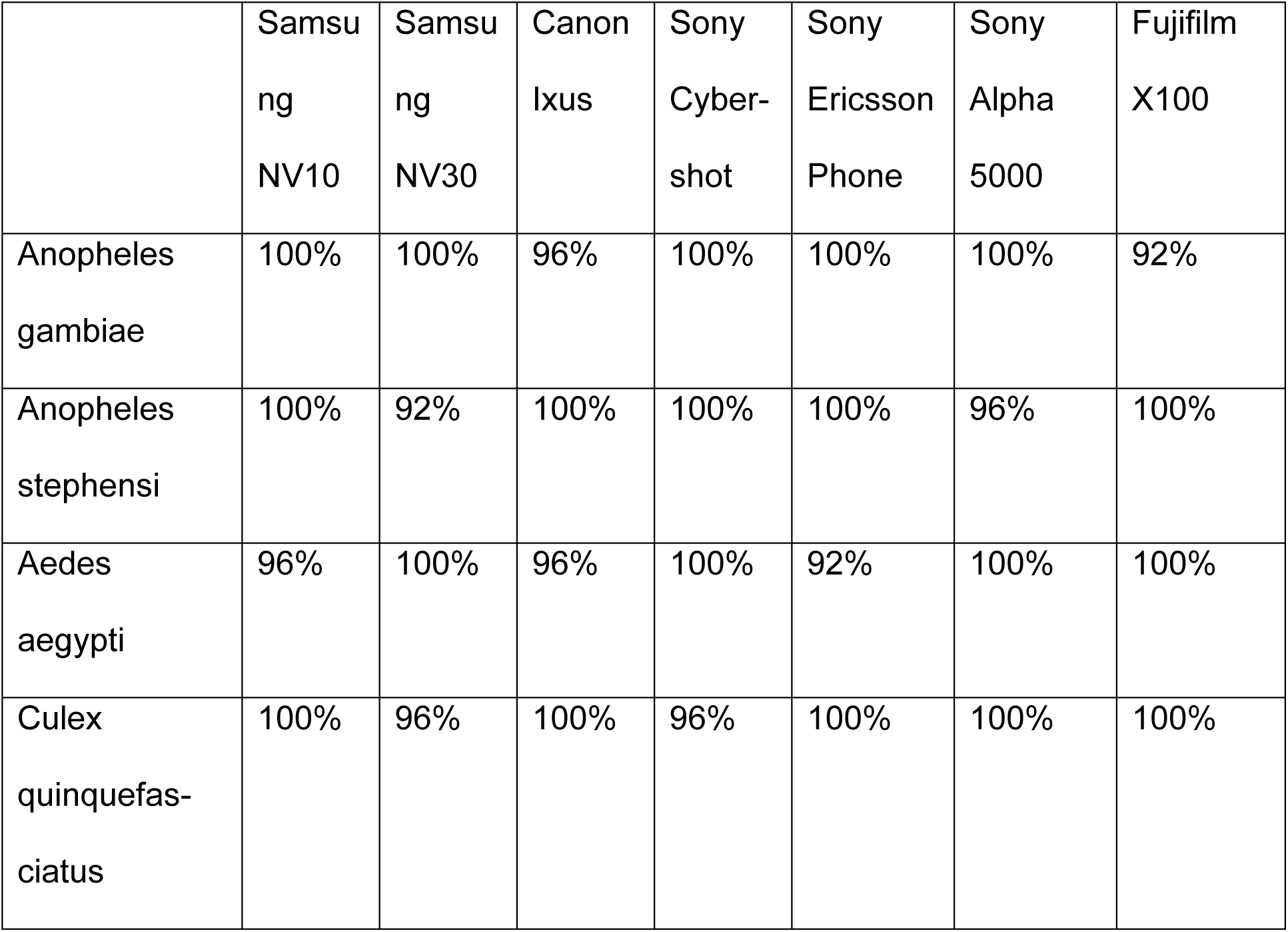
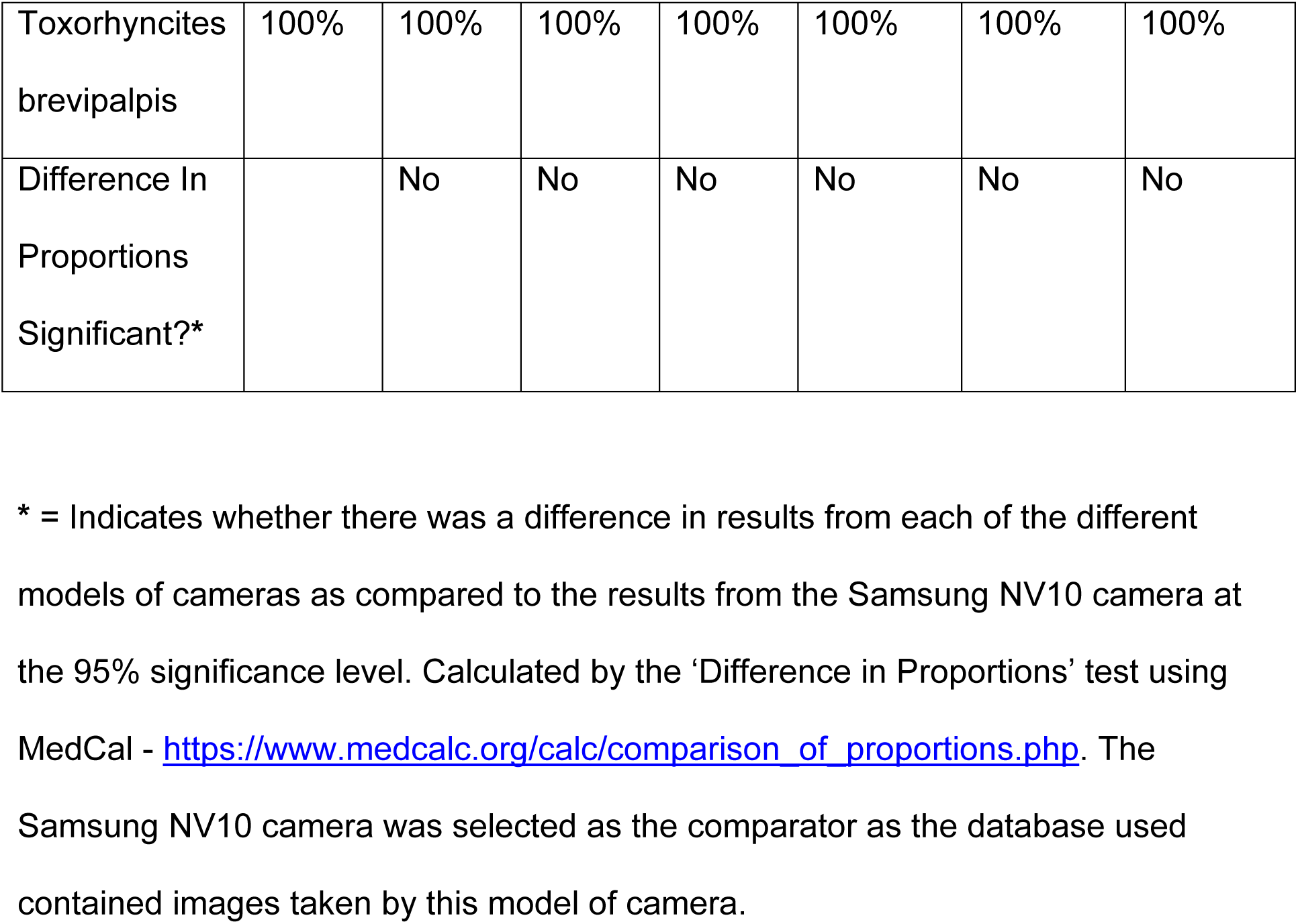
Percentage of correctly identified species ranked 1, using different models of cameras.

### Identifying Wild Caught and Laboratory Reared Species, Using One Male and One Female Wing in the Database

Using one male and one female wing image in the database, 34 different species of mosquitoes, two sub species of bees and two sub species of bumblebee wing images were tested and correctly identified rank 1 and rank 2 identifications noted. The numbers and sexes of each species available for testing varied and is given next to the species name in Table 5. Where only one sex was available, only one image was used in the database of the same sex being tested.

**Table 5:**
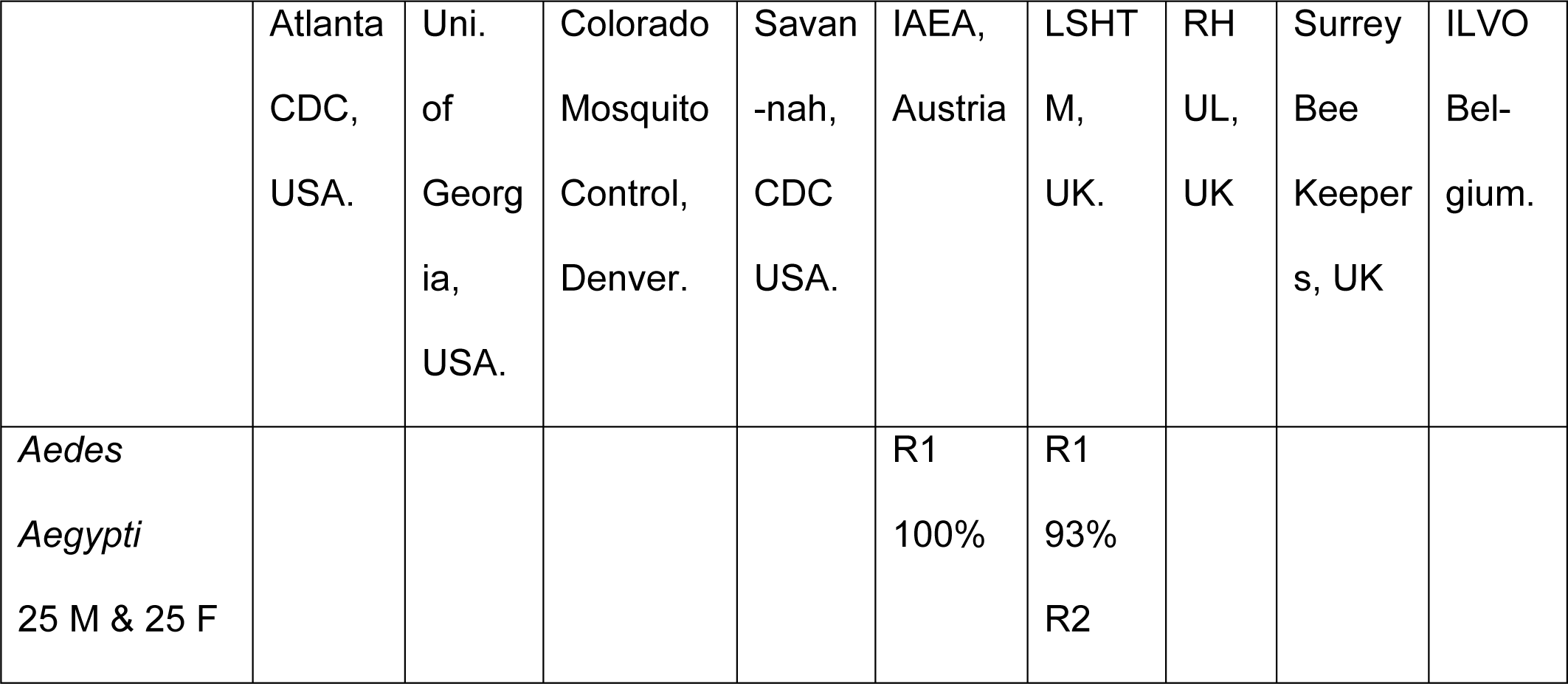
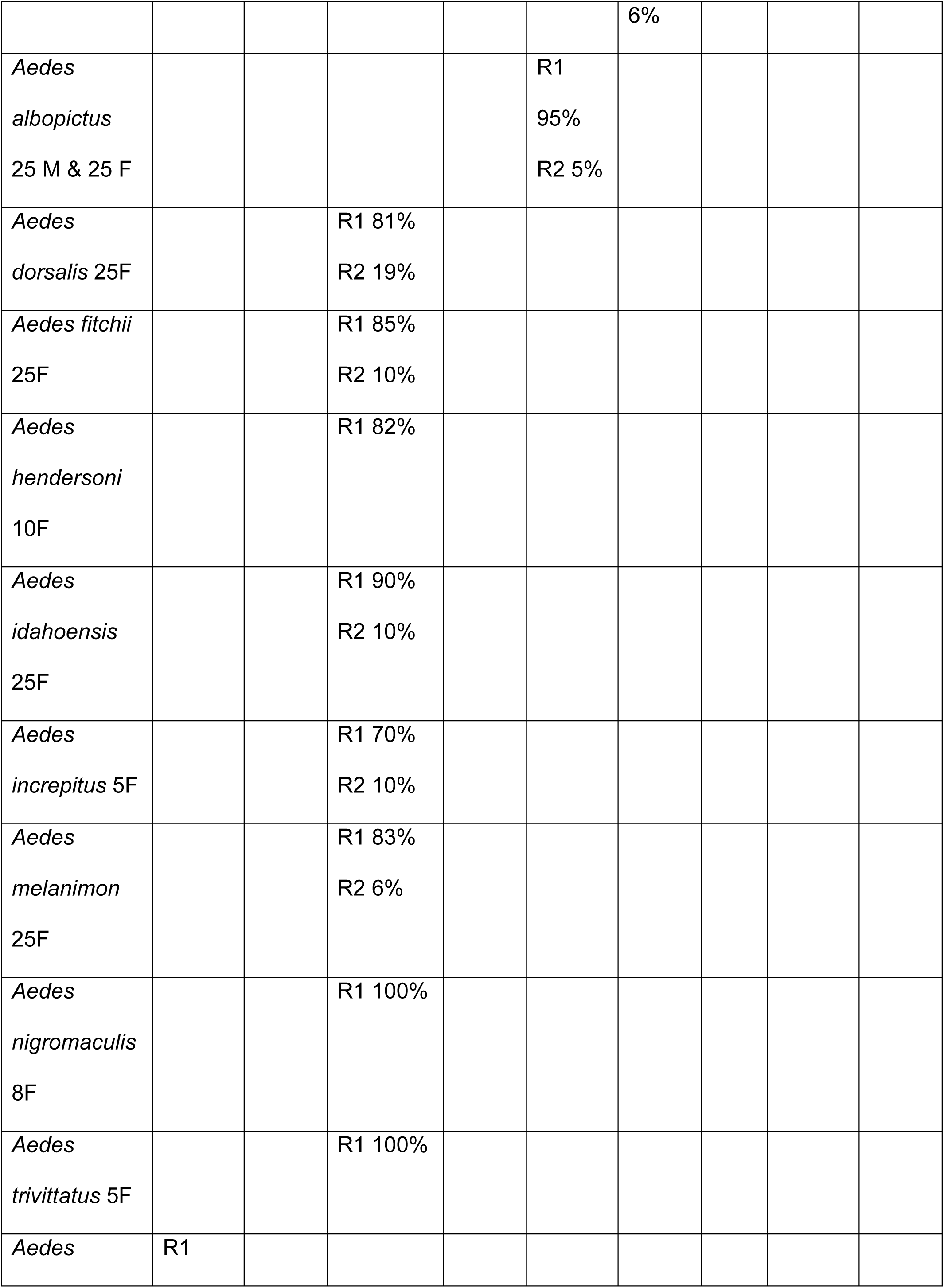
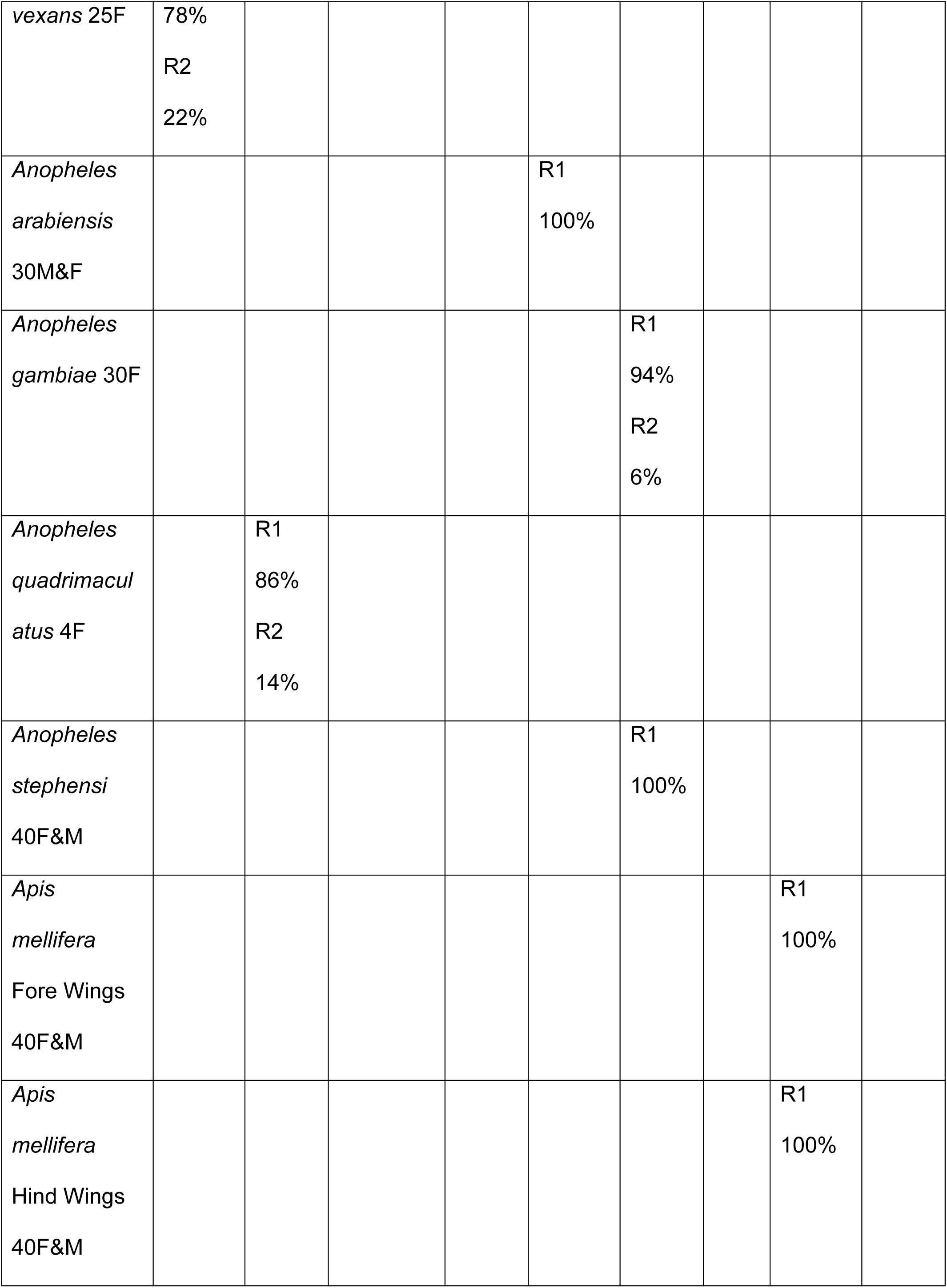
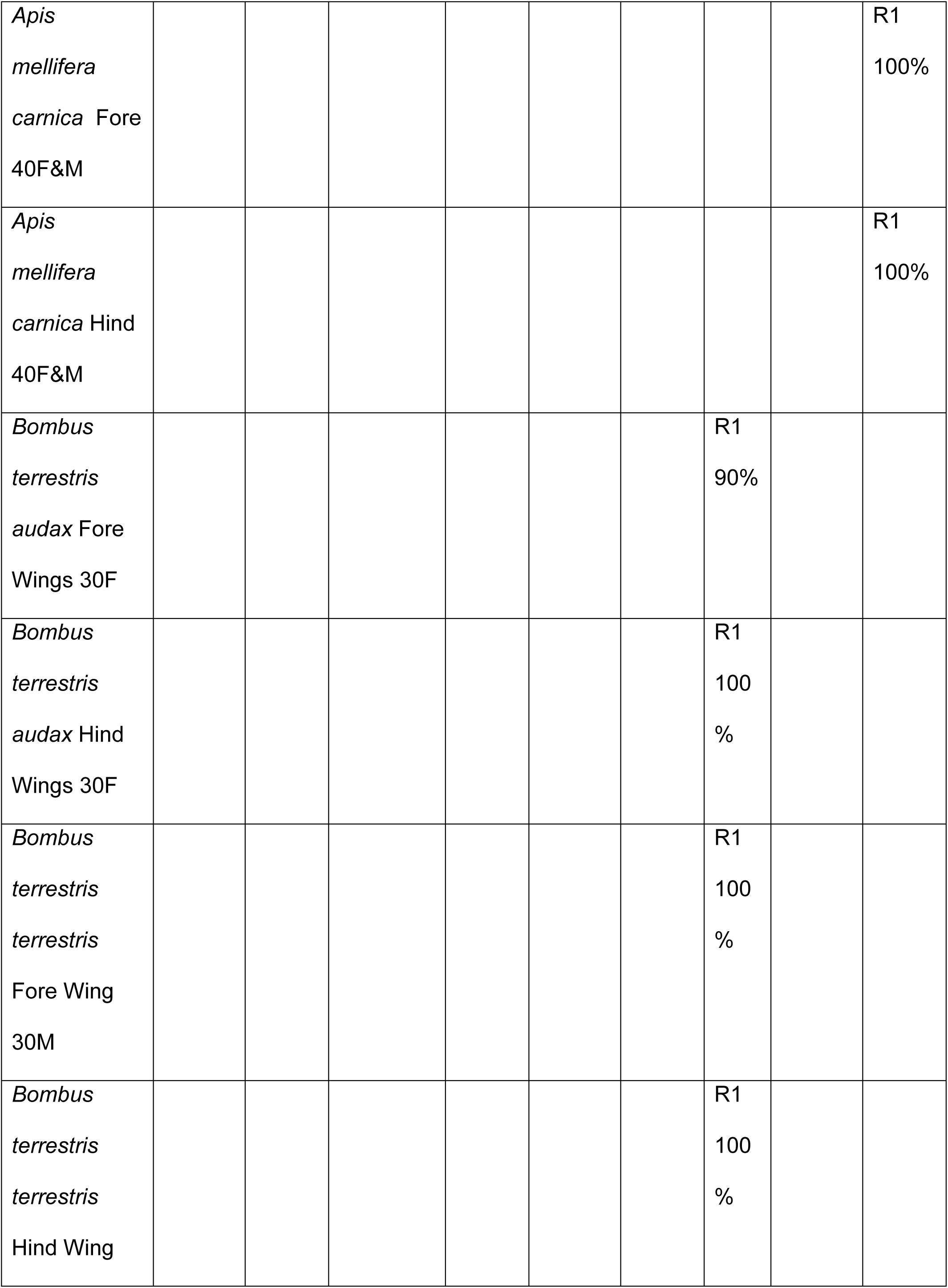
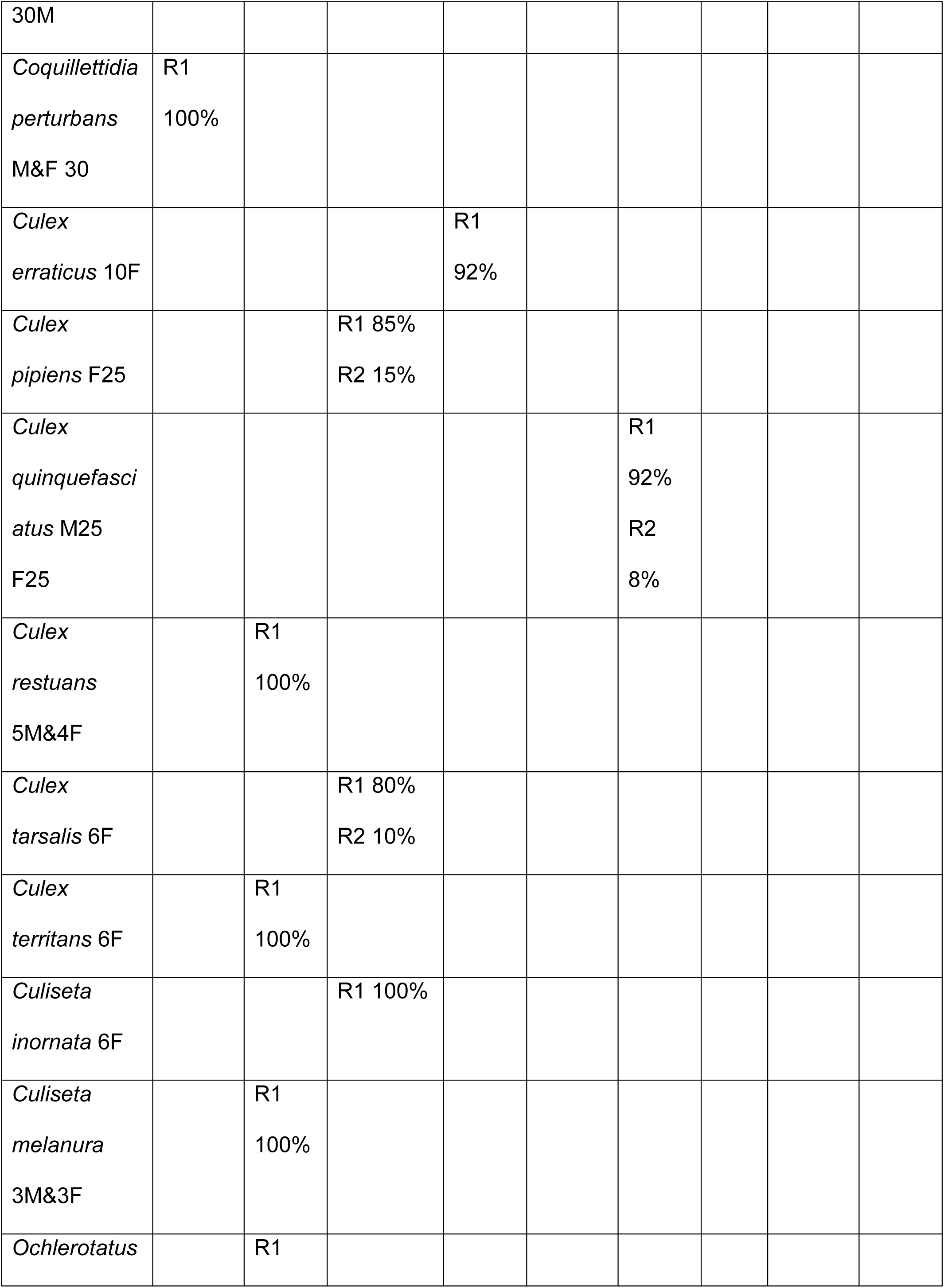
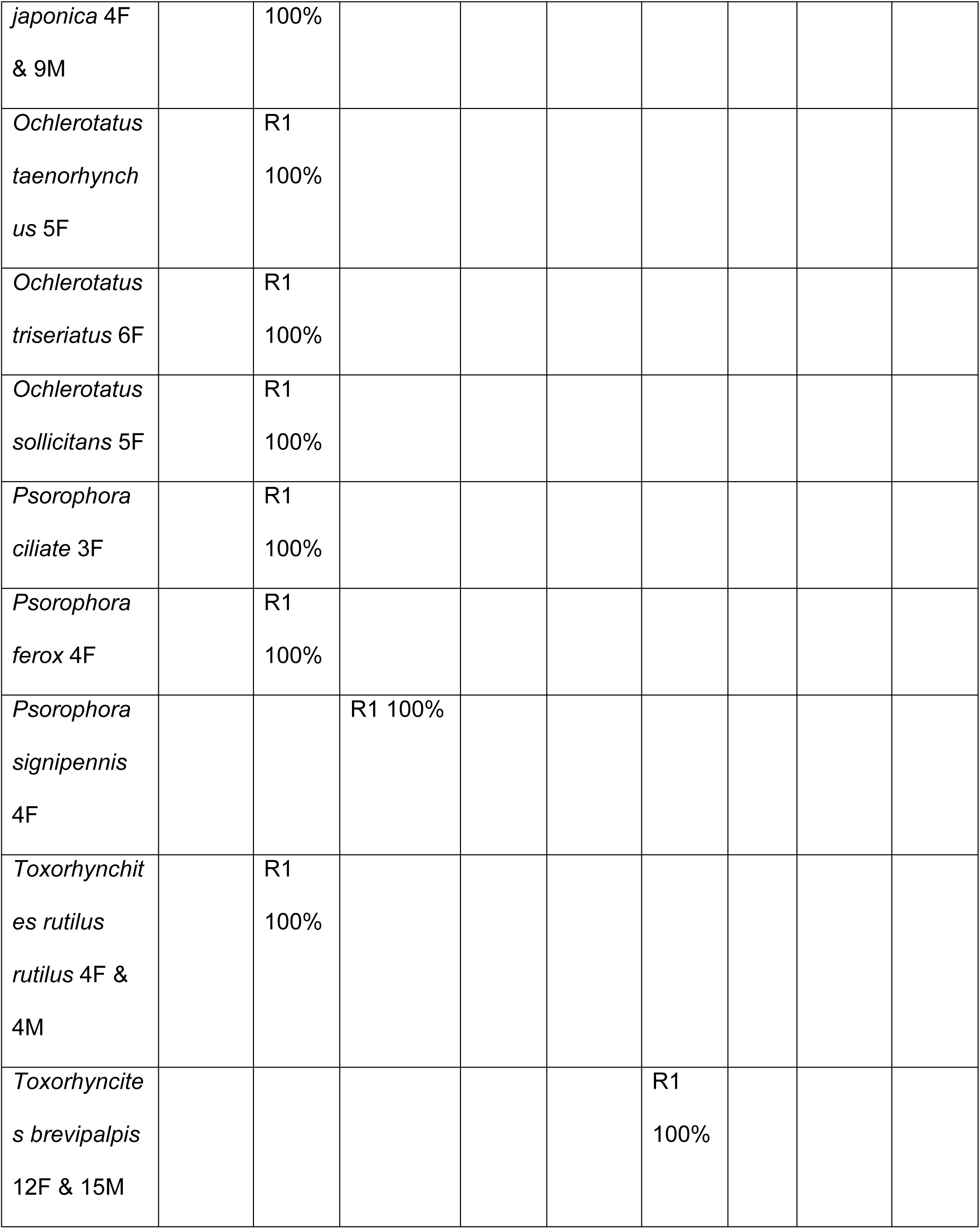
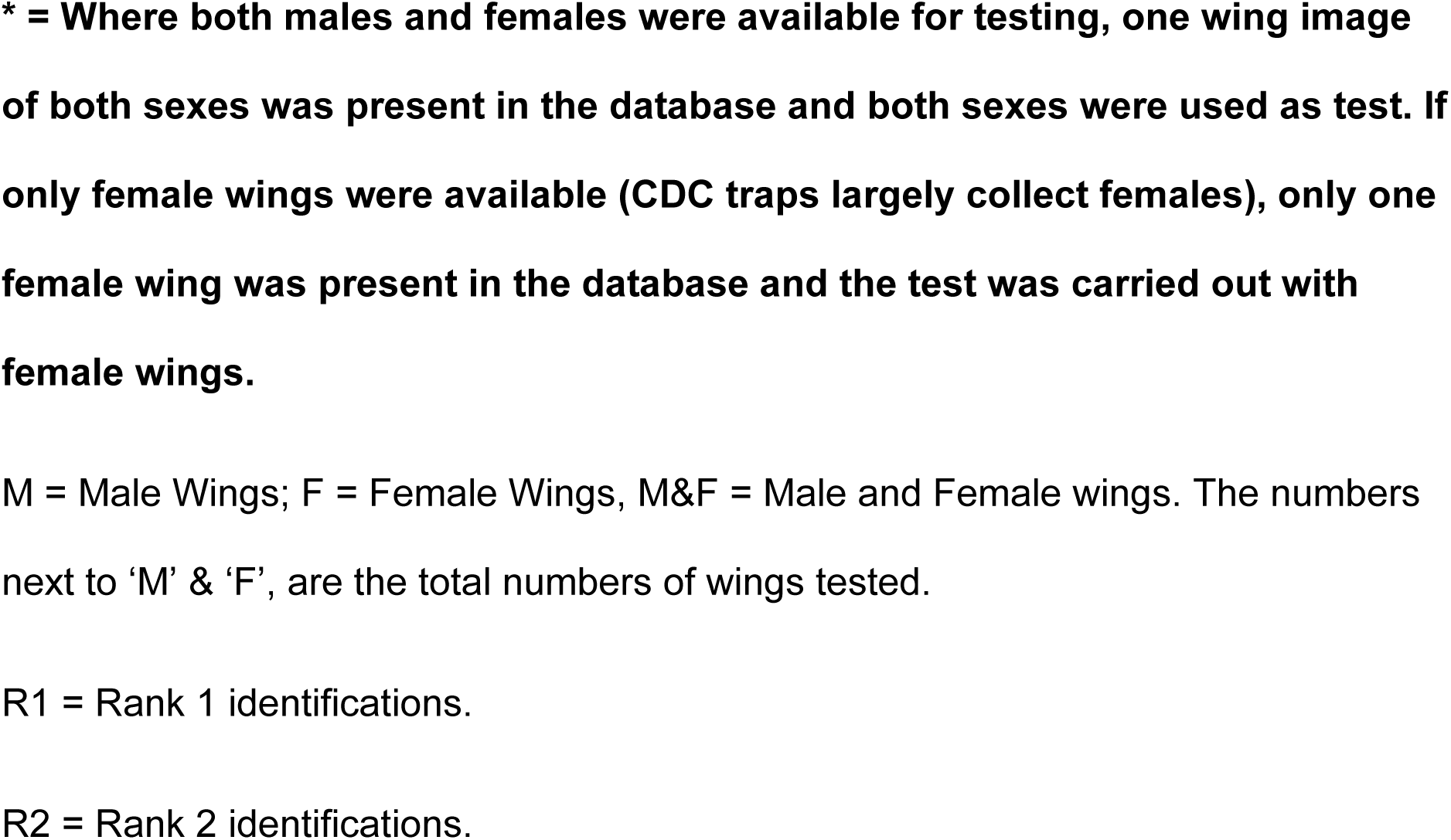
Identification of wild caught and laboratory reared species using one male and one female wing* in the database.

## Results

All of the species were wild, field caught specimens and identified by the respective donors. The available location data is presented. The LSHTM and the IAEA specimens were subsequently reared in their laboratories. The bee and bumblebees were from donor maintained colonies.

CDC = Centre for Disease Control, USA, Public Health Division.

IAEA = International Atomic Energy Agency, Insect Pest Control, Vienna, Austria.

ILVO = Instituut voor Landbouw-en Visserijonderzoek, Belgium.

LSHTM = London School of Hygiene and Tropical Medicine, UK.

RHUL = Royal Holloway University of London, UK.

1-DB = Total % correctly identified at rank 1 when using the database containing only one representative species and subspecies from the Diptera and Hymenoptera (includes fore and hind wings where appropriate).

2-DB = Total % correctly identified at rank 1 when using the database containing two representative species (one of each sex) of the Dipteran species and *A. mellifera* (includes fore and hind wings). N/A = Not applicable as only one sex was available in the case of Btt subspecies.

LDB and SDB = Total % correctly identified at rank 1 when using the Larger (20 of each species, 10 of each sex) and Smaller (10 of each species, five of each sex) databases.

‘T’ tests were carried out throughout using Graph Pad, QuickCalc software, http://www.graphpad.com/quickcalcs/ttest1.cfm.

A ‘t’ test of the LDB and SDB results indicated that the calculated value of ‘t’ was less than the critical ‘t’ value of 1.2394 (p= 0.05; df=20; SE= 1.98) and the ‘p’ value was more than 0.05, indicating that there was no difference between LDB and SDB.

A ‘t’ test of the 1-DB and LDB results indicated that the calculated value of ‘t’ was greater than the critical ‘t’ value of 4.018 (p= 0.05; df=20; SE= 6.334) and the ‘p’ value was less than 0.05, indicating that there was a significant difference between 1-DB and LDB.

A ‘t’ test of the 1-DB and SDB results indicated that the calculated value of ‘t’ was greater than the critical ‘t’ value of 3.547 (p= 0.05; df=20; SE= 6.664) and the ‘p’ value was less than 0.05, indicating that there was a significant difference between 1-DB and SDB.

**Up to Rank 5 Modal Values**: The calculated value of ‘t’ (1.7252) was smaller than the critical ‘t’ value of 2.086 and the p value (0.099) was greater than 0.05; df = 20; SE = 0.263; therefore there was no difference between the modes of the larger and smaller databases for up to rank 5 results of accurately identified species and sexes of wings.

**Up to Rank 10 Modal Values**: The calculated value of ‘t’ (3.7849) was greater than the critical ‘t’ value of 2.086 and the ‘p’ value (0.0012) was less than 0.05, df = 20, SE = 0.458; indicating that there was a significant difference between the databases and that when accurate species identifications are considered up to rank 10, the larger database results produced significantly larger numbers of correctly identified results.

The blue reference and the red landmark points have been exaggerated (made larger) for ease of viewing. Where the veins did not quite meet the edges of the wing as in the Hymenoptera, extrapolating the point still lead to accurate results, as long as it was uniformly observed in all images. I^3^S Classic, default setting, required a minimum of 12 points to be marked and allowed a maximum of 30 points that could be marked. The numbers of maximum and minimum points could be changed, however, the default setting was kept. Hence, extrapolation of points was required for hind wings to bring the marking up to the required 12 marked points.

## Discussion

A method of preparing and using insect wings for species identification in I^3^S was elucidated (Figures 2 and 3) and tested (Tables 1-5). I^3^S, initially developed to identify Individual whale sharks (Hartog and Reijns, 2013) was adapted and used for insect wing identification. The results indicate that when adapted, it can also be reliably used to aid the identification of insect wings, as images of different insect species were correctly identified with a high degree of accuracy, Tables 2, 3 and 5. Unlike CO1 (Ravela and Gamble, 2004), software which analyses the entire wing image, I^3^S only takes into account the markings made by the user (Figures 2 and 3). Any other diagnostic characteristic such as the shape of the scales, for example the diagnostic tear drop scales on *Coquillettidia perturbans,* or the patches of light and dark scales on *Anopheles* species, is not used by I^3^S. Future image recognition software should perhaps incorporate elements of both CO1 and I^3^S - whole image analysis as well as operator created markings, to advance species identification of insects using image recognition systems.

**Figure 3.**
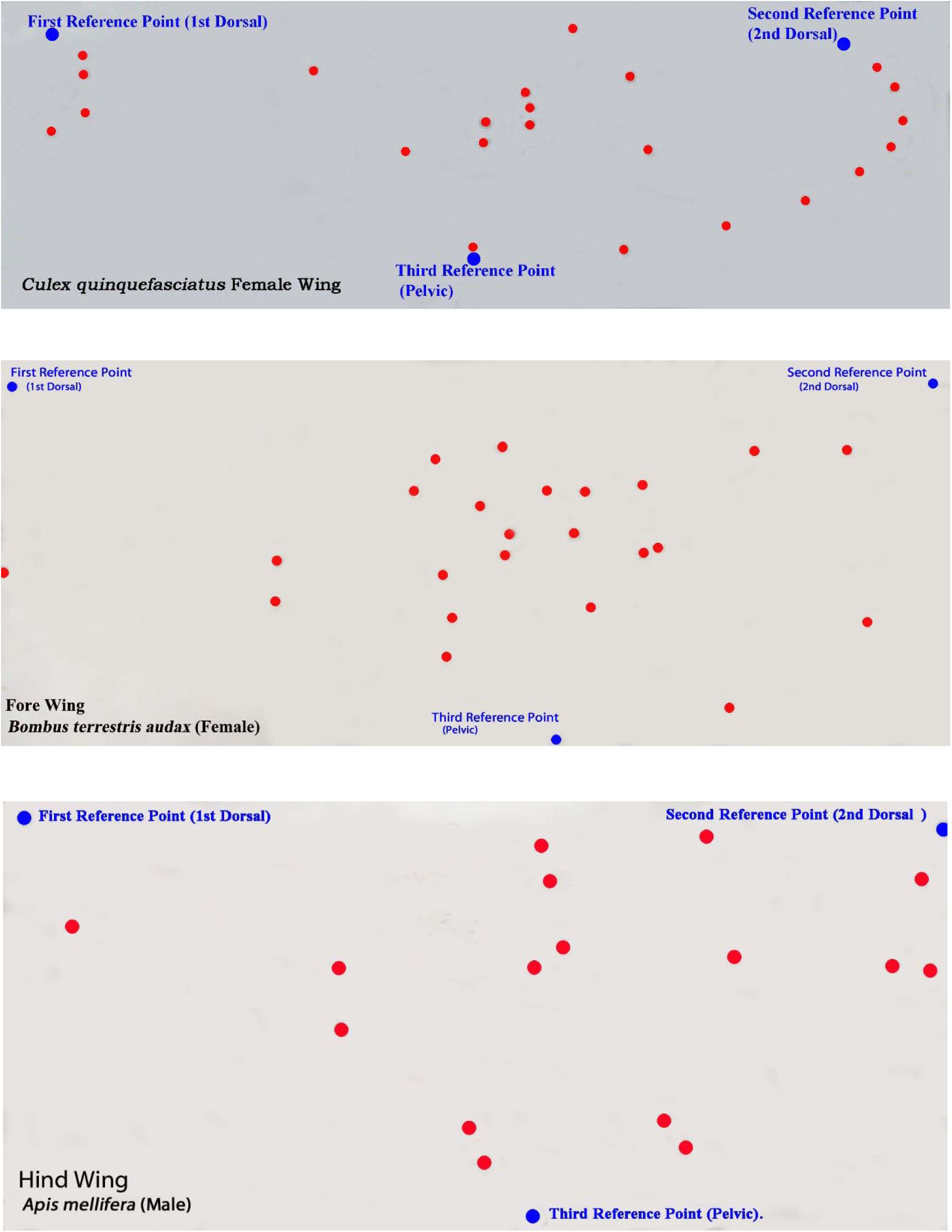
A visual depiction of the markings used by I^3^S to assess similarity with the images in the database.

The database was an important part of the identification process. Using just one image per species in the database was generally not sufficient to obtain high levels of accurate identification at rank 1 (Table 2). The only exception to this was the result for *Apis mellifera* where having just one specimen in the database was sufficient to obtain 100% accurate identification at rank 1 however small or large the database was, as its markings were sufficiently distinctive from all of the other wings in the database. The scores for *A. aegypti* and *T. brevipalpis* were also high when only one specimen was in the database (93% & 97%), probably because as with *A. mellifera*, their marking were very different to the other species in the database.

There was no difference between the results from the larger and smaller databases for rank 1 identifications when two images, one of each sex were present in the database. This was sufficient to ensure 100% accurate identification in rank 1 from all of the Dipteran and Hymenopteran species used - Table 2, indicating that I^3^S Classic was reliable and accurate software when at least two images, one of each sex, were present in the database. The necessity for using both sexes in the database was a result of the sexual dimorphism that exists between male and female wings in mosquitoes and other insect species (Virginio F, 2015) and the fact that I^3^S assess the proportional distance between the vein markings – this would be different if the wing shape and size was different between the sexes, however small that distance and shape was. In the majority of the Dipteran species tested, the tendency was to rank at 1, an image of a wing that was the same sex as that of the test but incorrect species, despite the fact that an image of the opposite sex of the correct species was present in the database. For example, using a database containing just one male *A. gambiae* wing, a test image of a female *A. gambiae* could result in a female *A. stephensi, or A. aegypti or C. quinquefasciatus* being ranked 1, even though a male *A. gambiae* was present in the database. This only occurred in databases that contained only one image of the test species (1-DB in Table 2). In the majority of the species tested, especially Dipteran species, images of both sexes were required in the database for accurate wing identification in I^3^S. This did not apply to the Hymenopteran species tested here. There were also no differences in rank 1 results when using larger or smaller databases (Table 2).

As all the database and test images were aligned the same way and marked consistently, another feature of the ranked results that could be assessed was how frequently the correct species was ranked from rank 1 to 5 and 1 to 10 (the ‘Modes’). The modes of correctly identified species (how frequently a particular species appeared within ranks) could be used to gauge the accuracy and similarity of retrieved results from ranks 1 to 5 and 1 to 10 (Table 2a). This indicated that although there was no difference in rank 1 and up to rank 5 identifications between the LDB and SDB (Table 2 & 2a), using a larger database produced larger numbers of accurate results up to rank 10 than a smaller database (Table 2a). However, this was to be expected as the SDB had five male and five female wings, rather than the 10 of each sex in the LDB. The SDB therefore only had the potential to bring up the first five of the same sex as the test when rank 1 to 10 was considered. Noting frequently identified species up to rank 5 and 10 can still be useful, especially where the correct species was not retrieved at rank 1, but was in rank 2, 3, 4, or 5. Although using one image of each sex produced high results, having more than one image of each sex in the database could add to certainty that the correct species was being retrieved if it appeared often enough up to rank 5 and 10 (the mode) in databases that contained more than 1 image of the species. Having five or 10 images of each species in the database and noting their retrieval in the first five or 10 ranks, is another way to add assurance.

An earlier study (Vyas-Patel et al, 2015) observed that if there was more than one wing specimen of a species in the database, it was ranked consecutively correctly further down the ranks. This feature was examined here. It was found that in the majority of cases, if there were five specimens in a database, all five were retrieved and ranked consecutively correctly. In no cases were only two retrieved consecutively correctly, there were at least three consecutively correctly ranked, Table 3 and associated Figure 1.

Using different camera models was not predicted to affect the results and Table 4 bore this out. This was because I^3^S works on the proportional distance between markings and this distance was never changed at any stage during the course of this study. Most cameras, whether highly sophisticated, like the Fujifilm X100, or a simple phone camera (Sony Ericsson), are capable of capturing the same, necessary detail of the veins, as it appears on the wings, in equal measure and usually with the same clarity, if the camera is used correctly. Only clear, sharply focused images were used. Furthermore, the smaller Dipteran wings were kept flat with a coverslip to prevent any folding of the wings. This was not required for the larger Hymenopteran wings. These results indicate that I^3^S can be used by the trained/citizen scientist without the need for an expensive camera, or by anyone with an ‘off the shelf’ camera and that using different models of camera would not affect the accuracy of the results, provided the images were in focus, aligned and marked the same way as the images in the database.

When different species from different locations/countries were considered and using a database that contained one image of each sex, it was found that there were high levels of accurate retrievals of the correct species (Table 5). In the few cases where the correct identification did not occur at rank 1, it was found at rank 2 indicating that when unsure of a test result, all of the ranked images should be examined, at least up to rank 5. It was not feasible to have more than one or two sample images in the database of each species/sex, as the numbers of species from some of the field collected specimens was low. Ideally, five and up to 10 images of each species and sex in the database would have added extra certainty to the results. With just one image of each sex, the average results from each donor species ranged from 89% to 100%. It was noticed that when the correct species occurred at rank 2, the rank 1 result always had the correct genus. This was particularly true of the *Aedes* species from Denver, where the genus (*Aedes*) would be correct at rank 1 and the species correct at rank 2. This indicates that the morphological differences between *Aedes* species was very small and that I^3^S was very accurate at ranking the correct genus at rank 1. Seldom was the genus incorrect at rank 1.

The separation between the major Dengue vectors, *A. aegypti* and *A. albopictus,* was high (93%, 95%) in keeping with the study by Henry et al (2010), where Geometric morphometric techniques were used to differentiate between the two species obtained from different locations.

The separation of sibling species (*Apis mellifera* and *Apis mellifera carnica; Bombus terrestris terrestris* (*Btt*) and *Bombus terrestris audax (Bta)*; *Anopheles gambiae* and *Anopheles arabiensis*) using image recognition, shown in Table 5, can undoubtedly be of great use in separating/differentiating between sibling species. Previous studies proved that morphological separation can be used to separate *A. mellifera* from *A. mellifera carnica* using ‘Geometric Morphometric techniques’ and that this separation tallied strongly with microsatellite separation (Oleksa and Tofilski, 2015). The results presented here lend further proof to the findings by Oleksa and Tofilski (2015) - I^3^S separated and identified both *Apis* sub species with 100% accuracy. The finding that other sub-species - *Btt and Bta; A. gambiae* and *A. arabiensis* could also be separated and identified accurately at high levels is also encouraging. In this study, the different sub-species had already been separated using molecular techniques. However, if an unknown species collected from the wild were to be used in I^3^S, then conclusive evidence of a given sibling species should only be made via rigorous molecular and morphological testing. It is only when very large numbers of wild species (suspected to be sibling/sub-species), have been rigorously tested using image recognition and substantiated with DNA molecular profiling, can the final sibling/sub-species diagnosis be made. The method and the results in Table 5, provides for a morphological separation of sibling species using I^3^S image recognition that was not possible by eye previously. Furthermore, the sibling species came from different locations and laboratories, very far apart and had been originally verified using molecular techniques, making this feature of the results worthy of further scrutiny and study. The results from the current study certainly suggests strongly that morphological separation using I^3^S image recognition is in keeping with their molecular separation.

These are promising results for the future use of image recognition software. However, the high rate of accurately identified results in this study are due to the fact that great care had been taken on a number of levels as the primary aim of the study was to test the software, not the principals of using image recognition for insect species identification. Images of all of the species used as test had copies in the database, without which it would have been impossible to test the software. Unless the software was used with care, keeping a number of points stated below in mind, high levels of accurate species identification could be compromised.

Image recognition software for insect species identification should always be used intelligently, with care and bearing a number of important points in mind. The first of these is that the accuracy of any identification could only be as good as the images in the database. Both test and database images should be clear, adequately focused, marked consistently and aligned the same way. Secondly, it should be kept in mind that if an image of the test was not in the database, only an image that next resembled the test would be pulled out of the database and ranked at 1. Furthermore if there were five other images of the same species that closely resembled the test (but were not of the same species), then these images would also be pulled out of the database and more than likely ranked consecutively from rank 1 to 5. The best way (amongst others) to counteract this, would be to not rely solely on one feature of the identification process, i.e. the wings. Features of the insect anatomy other than the wings should also be considered, as is the norm when arriving at identifications using traditional taxonomic keys, when totally unknown species are used as test. Databases of a host of other features - markings on the thorax or legs, the shape of the wing scales or patterns of light and dark patches on the wings for example, can be created and used to confirm or refute an identification. A study of the scores for each database should be carried out, to aid the identification process. The scores will vary for every new database used, but assessing the scores for totally alien and closely resembling images for each new database can give some indication as to how closely a test image resembled those from the ranked results (Vyas-Patel et al, 2015).

This does not mean that databases containing only one feature of the insect anatomy have no value; that would miss the point of this study which was to test the software to see how accurate it was in retrieving known species that had representative copies in the database. If confirmation is required of an unknown wing image, suspected to be of a particular species, then running a wing image through a software programme that contained wing images of the suspected species in its database, can either confirm or refute previous species identifications. It can also aid the taxonomic process where identification is carried out using traditional keys, to confirm or refute the identification at any stage of the traditional identification process. As long as the images used were focused and clear, had been aligned and marked consistently in the same way as the images in the database, the contents of which were known, then image recognition systems such as I^3^S can play a very useful role in insect species identification. Recent studies have shown that using semi-automated morphometric techniques, citizen scientists could indeed use wing morphology with great accuracy to distinguish between different species of invasive and economically important wood borers and had the potential to do so for corn borers (Goczał J 2016; Ricciuti E 2015; Przybylowicz L et al, 2015). I^3^S should prove just as amenable for use by citizen scientists and farmers whose livelihood and crop production capability depends on accurate pest species identification. As technology using software is a fast moving phenomenon, image recognition of insect species can only develop further. If image recognition systems retained accuracy as it developed, it should become commonplace and a very useful tool for both the trained and citizen scientist, in fact, anyone that needed to know which species of insect they were dealing with.

## Conclusions

- A method of preparing and testing I^3^S for the identification of insect species using their wings was described and tested. The results indicated that I^3^S can be reliable and accurate software to use in the identification of insect species using wing images.
- Only similarly aligned and marked, clear, focused images should be used, in both the databases and test. The contents of the database should be known.
- Using just one wing image of each sex in the database was enough to obtain high levels of accurate identifications. Using five or 10 images of each species and sex in the database and noting if they were ranked early on in the database could also aid the identification process.
- The camera model used did not affect the results provided that the proportional distance between the markings was never changed at any stage.
- It was possible to separate sibling species using I^3^S, but conclusive proof for the occurrence of any wild caught sibling species should be based on molecular as well as morphological testing of the wings in I^3^S.

## Acknowledgements

This study would not have been possible without the generous donation of insect specimens from The London School of Hygiene and Tropical Medicine London, UK (Miss Shahida Begum and Dr Mark Rowland); Royal Holloway College University of London, Surrey, UK (Dr Gemma Baron and Prof Mark Brown); Surrey Beekeeper’s Association, UK (Mr Peter Bowbrick), Centre for Disease Control (CDC) Atlanta, US (Dr Rosemarie Kelly); Colorado Mosquito Control Denver, US (Dr Michael Weissman); University of Georgia Atlanta, US (Dr Elmer Grey); International Atomic Energy Agency, Austria (Dr Jorge Hendrichs, Dr Jeremie Gilles, Mr David Almanar); Chatham County, Centre for Disease Control (CDC), Savannah US (Dr Laura FAW Peaty, Dr Robert A Moulis); Instituut voor Landbouw-en Visserijonderzoek, (ILVO) Belgium (Dr Wim Reybroeck). The study would also not have been possible without dialogue with the creators of the freely available software I^3^S, Mr Jurgen den Hartog and Miss Renate Reijns. Everyone’s time, interest, help and discussion is gratefully acknowledged.

